## Supplementary Material and Methods for "Oscillations support short latency co-firing of neurons during human episodic memory formation"

**This PDF file includes:**

Materials and Methods
Supplementary Text
Figs. S1 to S11
Table S1
Supplementary References

### Supplementary Text

#### Firing Rates

During encoding, trials which resulted in a complete memory (hits) showed a sustained increase in firing rates compared to trials resulting in incomplete or no memory (misses; see Figure S2). This "subsequent memory effect" (SME) started approximately 1 second after the onset of the associative stimuli. A two-way repeated measurements ANOVA with the factors Memory (Hits vs Miss) and Time (Cue, Association Stimulus, Response) revealed a significant interaction ($F_{2,434}=7.16$; $p<0.001$). This effect was due to hits showing significantly higher firing rates than misses in the Response period (3-5 seconds; $t_{217}=3.79$; $p<0.0001$). No significant differences were observed during the Cue period (0-2 seconds; $t_{217}=1.89$; $p>0.05$), or during the Association Stimulus onset (2-3 seconds; $t_{217}=0.95$; $p>0.3$). Significant SMEs in the Response period were also obtained in more conservative statistical approaches, where sessions were used as random variable ($t_{35}=2.25$; $p<0.05$).

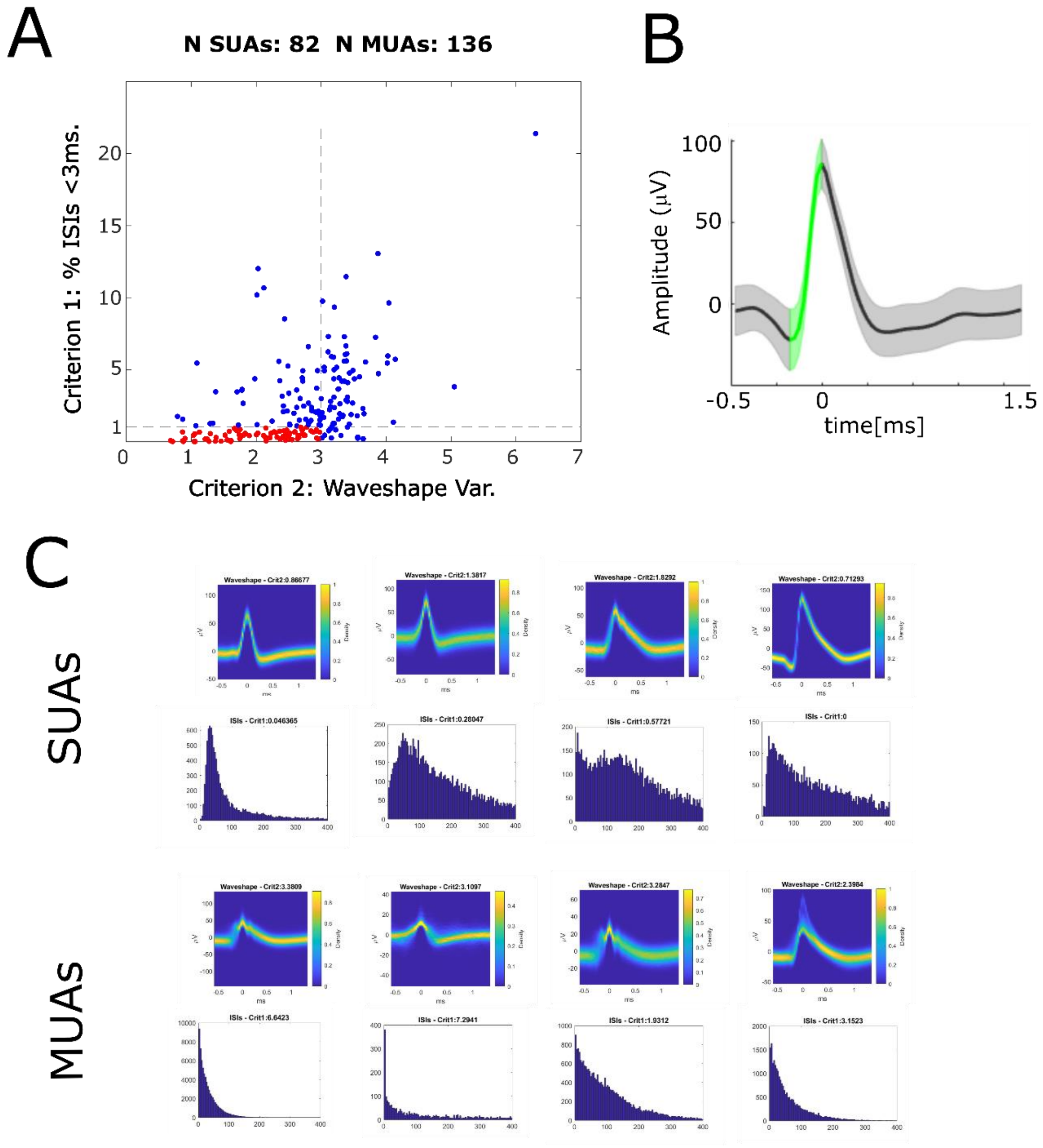


**Fig. S1. Automatic classification of Single- and Multi-Units according to Tankus et al. (*1*).** A) A scatter plot shows the distribution of the two criteria according to which neurons were classified into single-units and multi-units. Criterion 1 (y-axis) is the percentage of ISIs < 3ms. Criterion 2 (x-axis) is the variability of the spike waveshape in the rise time window. If a given unit shows less than 1% of ISIs < 3ms, and low variability of waveshapes (<3) then is labelled a single-unit (red dots), otherwise it is labelled a multi-unit (blue dots). B) The variability of the spike waveshape in the rise time window is shown for one example single-unit. The green shaded area highlights the rise time window which starts at the maximum curvature pre-peak, and ends at the peak. Waveshape variability is computed dividing the summed standard deviation in the rise time window by the rise height (i.e. peak-to-trough difference). C) Waveshapes and ISIs are plotted for 4 example SUAs (top row) and 4 example MUAs (bottom row).

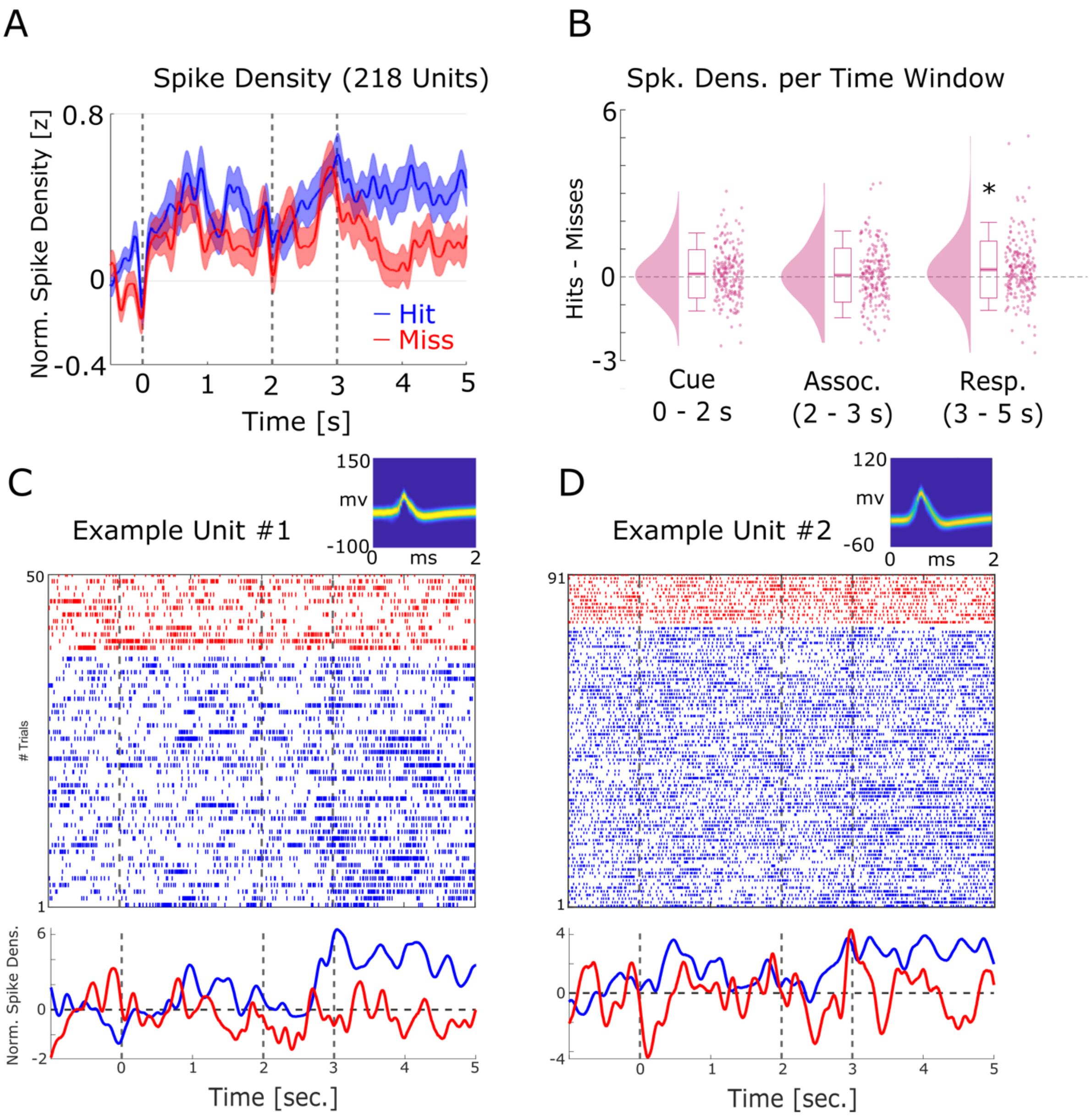


**Fig. S2. Firing rate effects during memory encoding.**
(A) Averaged normalized population spike densities are plotted for hits and misses. Hits show a sustained increase in firing after 3 seconds compared to misses. Shaded areas indicate standard error of the mean. (B) The difference between hits and misses is shown for the three time windows of interest. A significant increase for hits > misses was only observed for the response time period. (C) and (D) Two example single units are shown. Wave shapes on the top right are plotted by means of 2D histograms (see (*2*)). The plots beneath the raster plots show the normalized spike densities for hits (blue) and misses (red).

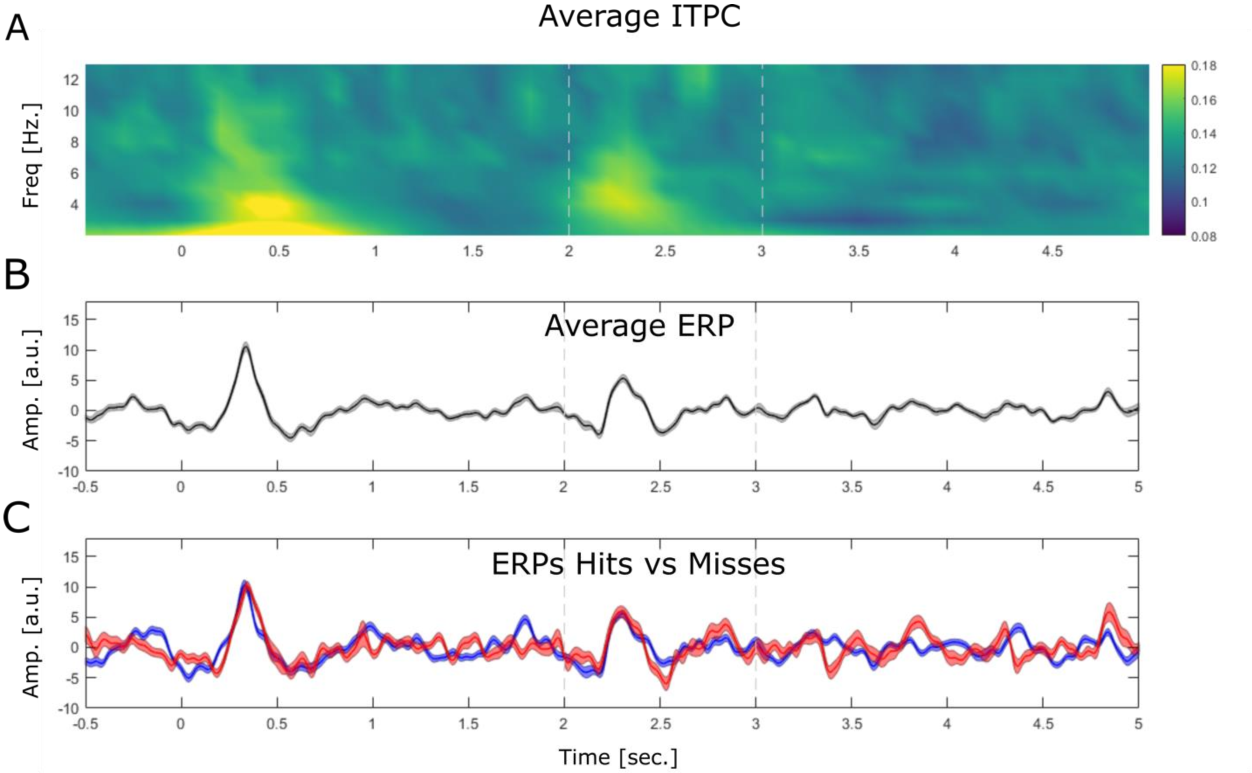


**Fig. S3. Stimulus evoked LFP activity is shown by means of inter-trial phase coherence (ITPC) and event related potentials (ERPs).**

(A) ITPC is plotted by means of phase locking value (PLV). The data shows a robust evoked response at the onset of the cue stimulus (0 sec.) and the onset of the association stimulus (2 sec.) albeit the latter appears to be slightly weaker. (B) The ERP is shown averaged across all trials, electrodes and sessions. (C) The ERP is shown for hits (blue) and misses (red). Note that no differences between ERP components between hits and misses is observed.

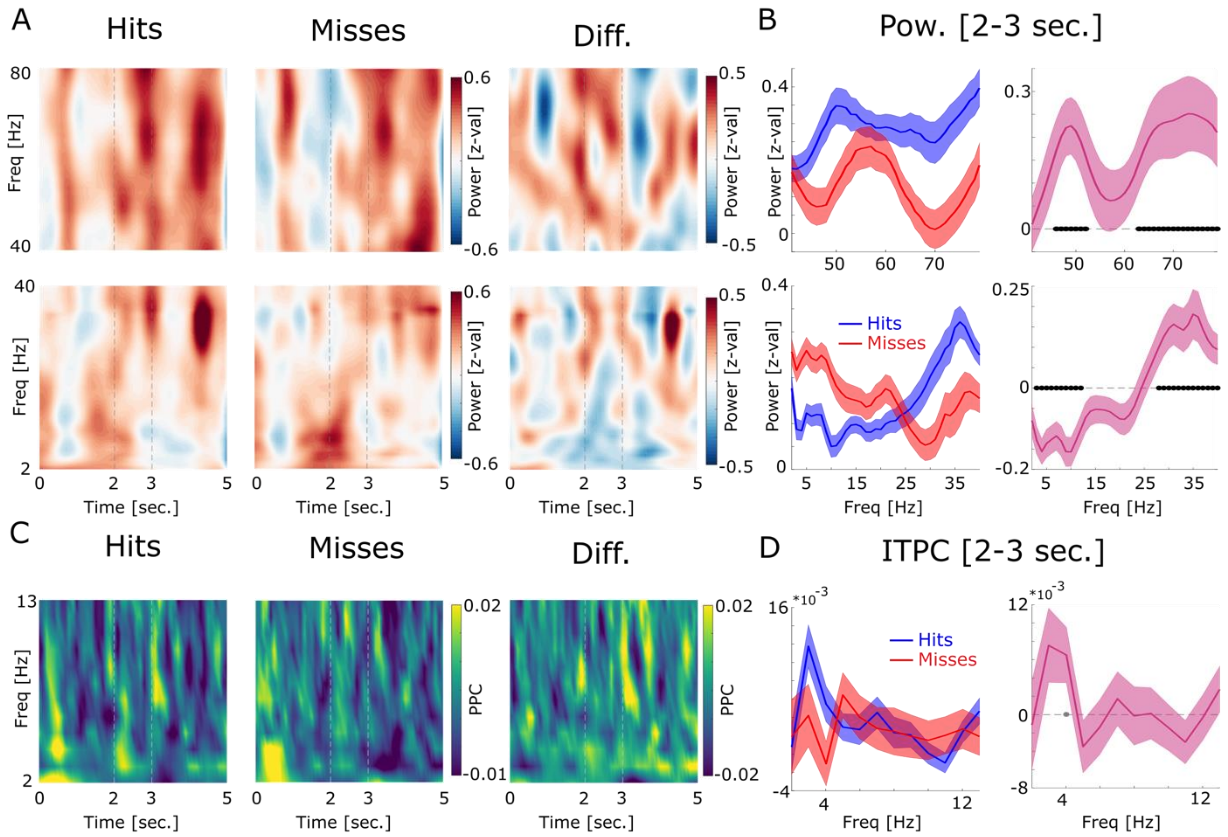


**Fig. S4. LFP power and intertrial phase coherence (ITPC) results are shown.**
(A) Power for high (top) and low frequencies are plotted for hits, misses and the difference. Time axis indicates time from cue onset. The dashed lines indicate the time window of association period. (B) Power during the association period is shown for hits (blue) and misses (red) and the difference (pink). Low frequencies show decreased power for later remembered associations, whereas high frequencies show increased power for later remembered associations which is a typically observed pattern (*3*). Concerning the higher frequency range, hits show increased power in a slow (45 – 50 Hz) and a fast gamma band (65 – 80 Hz) compared to misses. Filled black circles indicate $p<0.05$, FDR-corrected. (C) ITPC results are shown by means of pairwise phase consistency (PPC) for hits, misses and the difference. Increased phase concentration across trials is observed for the low theta frequency band at the onset of the cue stimulus, as would be expected, and somewhat weaker at the onset of the association stimulus (cf. Fig. S2). (D) PPC results are shown for the association time window (2-3 sec.) for hits, misses and the difference. No significant difference between hits and misses was observed, albeit hits showed a trend for increased phase coherence at 4 Hz compared to misses ($p<0.05$; uncorrected; grey filled circle).

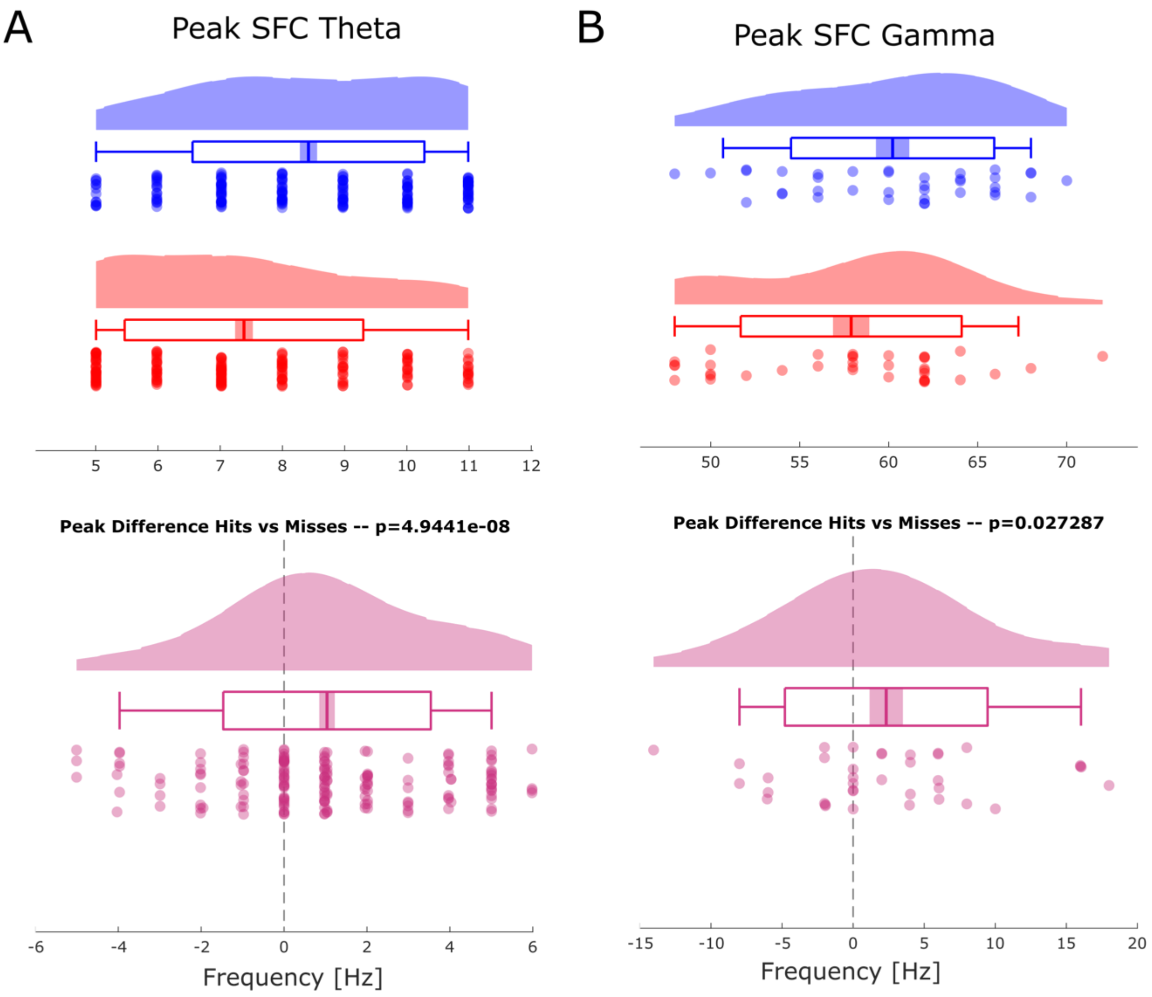


**Fig. S5. Selection bias control analysis**
Results of a control analysis are shown to rule out a possible bias on the Spike-Field Coupling results due to unbalanced trial numbers. Both control analysis replicated the original results showing faster theta frequencies for hits (blue) compared to misses (red, compare Figure 3 in main text), and faster gamma frequencies for hits compared to misses (compare Figure 2 in main text). (A) The results of the control analysis are shown for theta. (B) The results of the control analysis are shown for gamma.

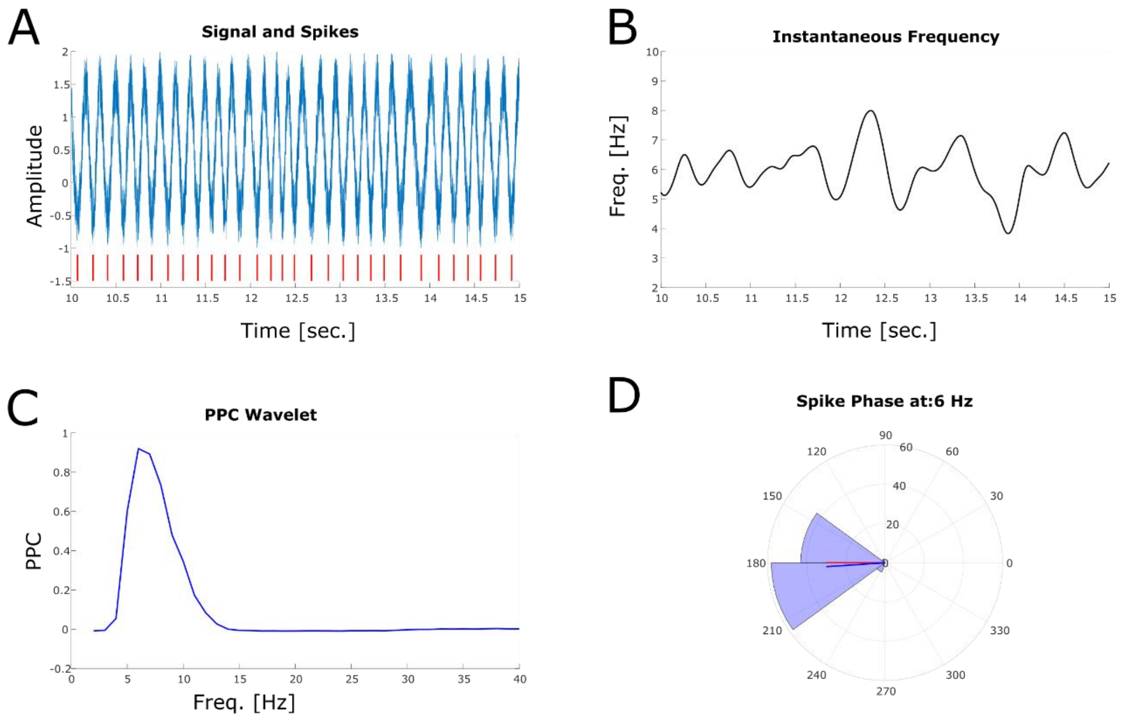


**Fig. S6. Simulation of the effects of a non-stationary oscillator on Wavelet analysis.** A) A signal is simulated that randomaly transitions between slower and faster frequencies with a mean frequency of 6 Hz (range: 3.5 – 9 Hz). White noise is added to the signal. Spikes are shown on the bottom (red ticks). Spikes are locked to the trough. An epoch of 42 seconds has been simulated but here only 5 seconds are shown. B) Instantaneous frequency is plotted (derived from the simulated signal before adding noise using 'instfreq' in Matlab). Note the strong non-stationarities in frequency. C) PPC spectrum shows a clear peak at the true mean frequency of 6 Hz. D) The phase histogram shows the phase derived from Wavelet analysis at 6 Hz. Mean angle of the spike phase is shown in blue, the ground truth phase angle is shown in red.

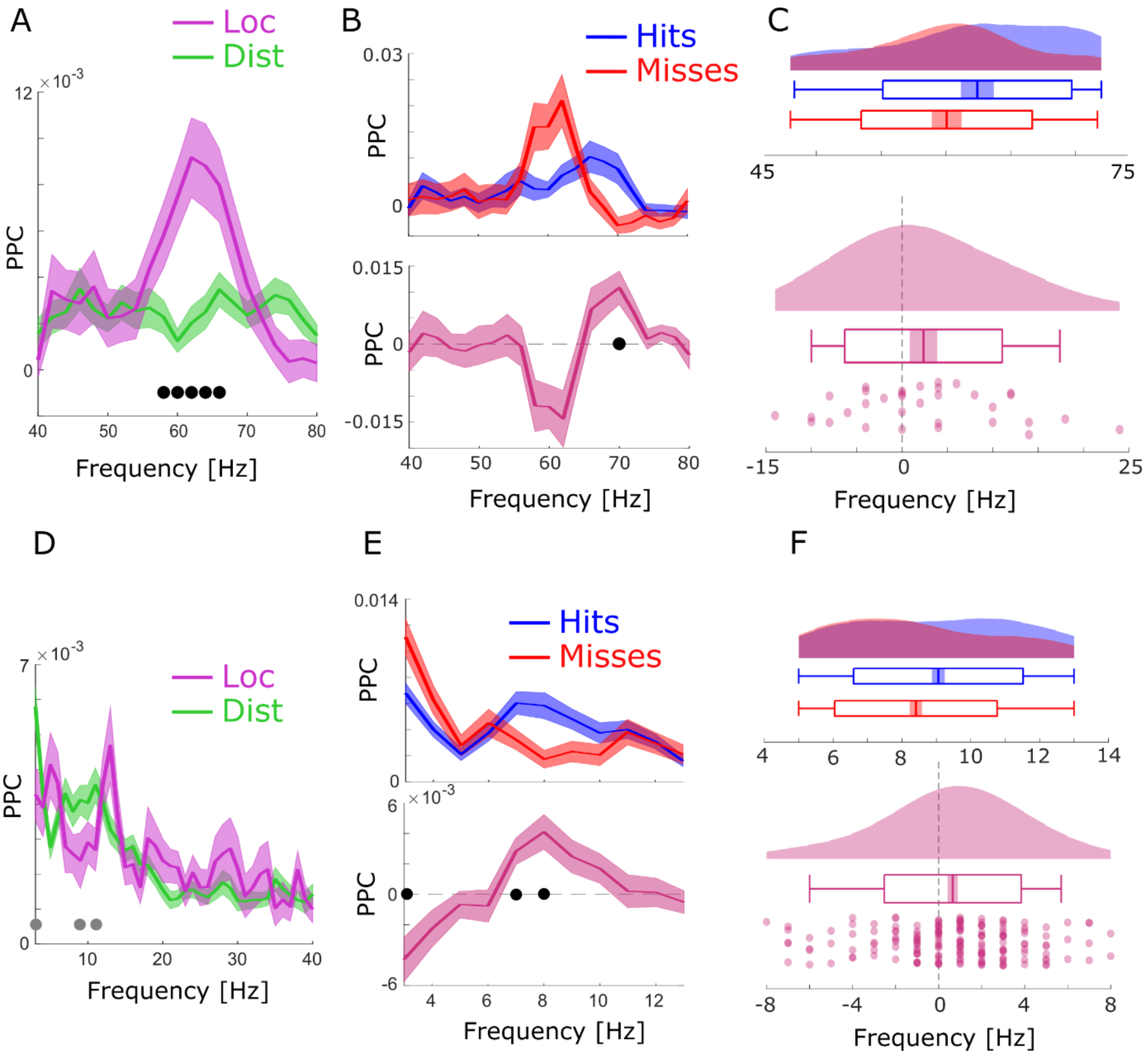


**Fig S7. Spike-LFP coupling results obtained with bandpass filtering and Hilbert transformation.** A) Spike-LFP coupling for the high frequency range for local (pink) and distal (green) pairs is shown. Local spike-LFP pairs show higher phase coupling compared to distal pairs in the high gamma range (black dots; $p_{corr}<0.05$). B) Top: Spike-LFP coupling is shown for the gamma frequency range for local pairs only for hits (blue) and misses (red). Bottom: The difference between hits and misses is shown. Black dots indicate $p_{corr}<0.05$ (FDR corrected). C) Gamma peak frequency in PPC across all local spike-field pairs is shown for hits and misses (top), and for the difference (hits-misses). Box plots indicate the same indices as in Fig 3D in the main manuscript. Hits trend towards faster gamma frequencies compared to misses ($t_{32}=1.56$; p=0.06). D-F) Same as in A-C for the low frequency range. Hits and misses are shown for distal spike-LFP pairs only. Hits demonstrate significantly higher theta frequencies compared to misses ($t_{175}=2.73$; p<0.005).

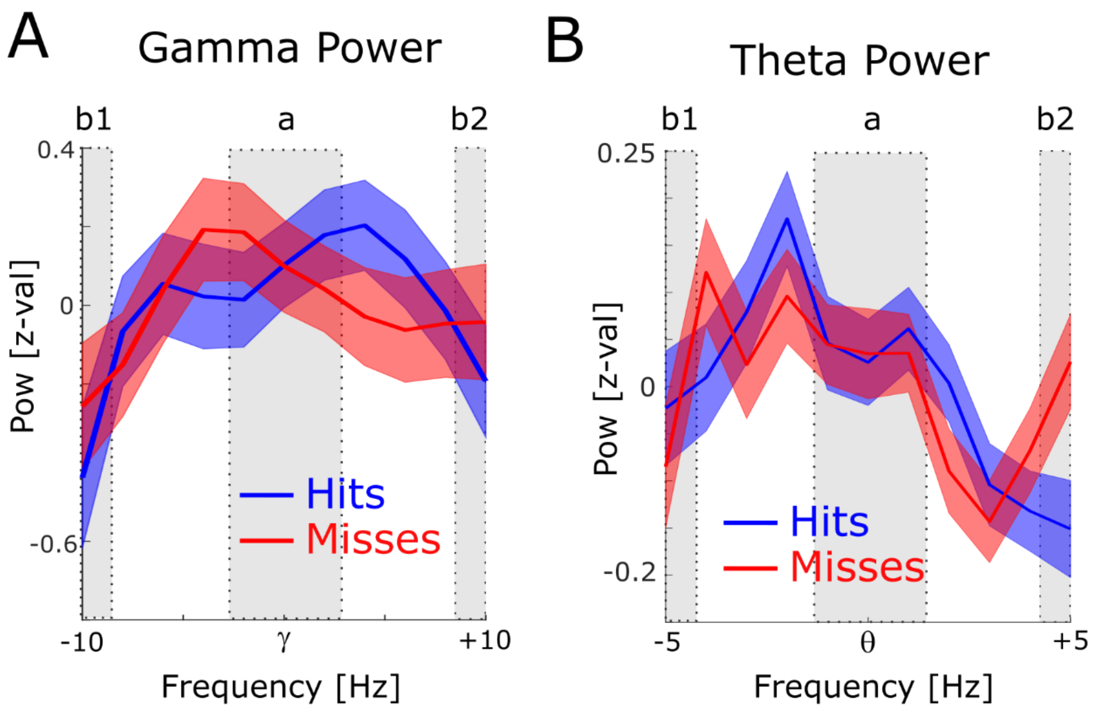


**Fig. S8. Power for phase providing theta and gamma frequencies and PPC for high and low power spikes.** A) The 1/f corrected power spectra are shown for hits (blue) and misses (red) centred on the peak gamma frequency of spike-LFP coupling. To demonstrate the existence of a meaningful signal in the phase providing frequency range (i.e. peak in the power spectrum) a paired samples t-test was calculated, where the power in the peak (a) was contrasted with the power at the edges (b1 and b2). Power values were averaged for hits and misses. For the gamma range, power at the peak was significantly higher compared to the power at the edges ($t_{52}$=2.38; $p<0.05$). B) Same as A but for the theta frequency range. Theta power at the peak was significantly higher compared to the power at the edges ($t_{374}$=2.44; $p<0.01$).

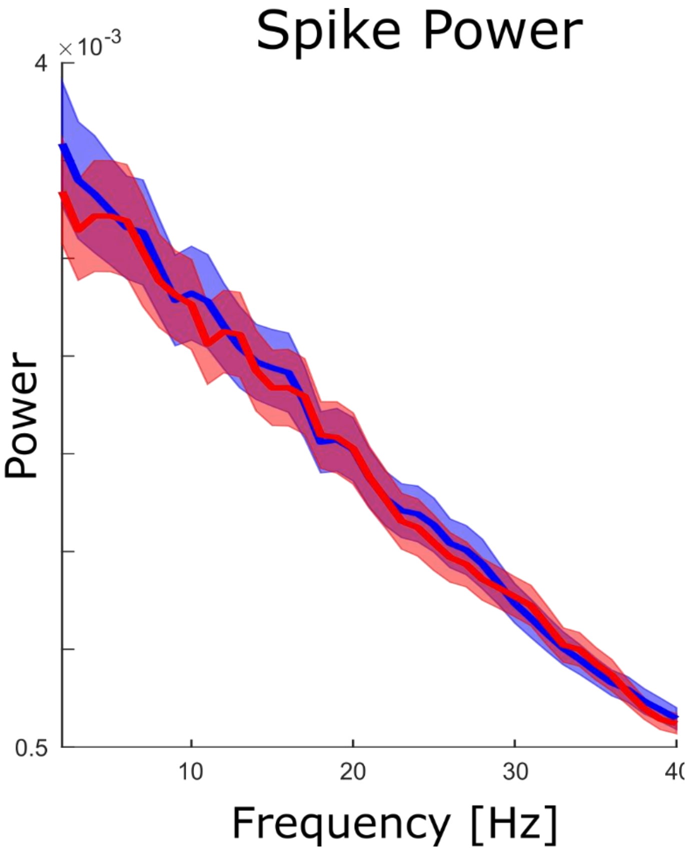


**Fig. S9. Spike Power Analysis.**
An FFT analysis is shown for continuous spike density time series to test whether spikes themselves showed clear peaks in the theta frequency range. No peaks were obtained neither for hits (blue) nor for misses (red), and no significant differences between hits and misses were observed ($p_{corr}>0.05$).

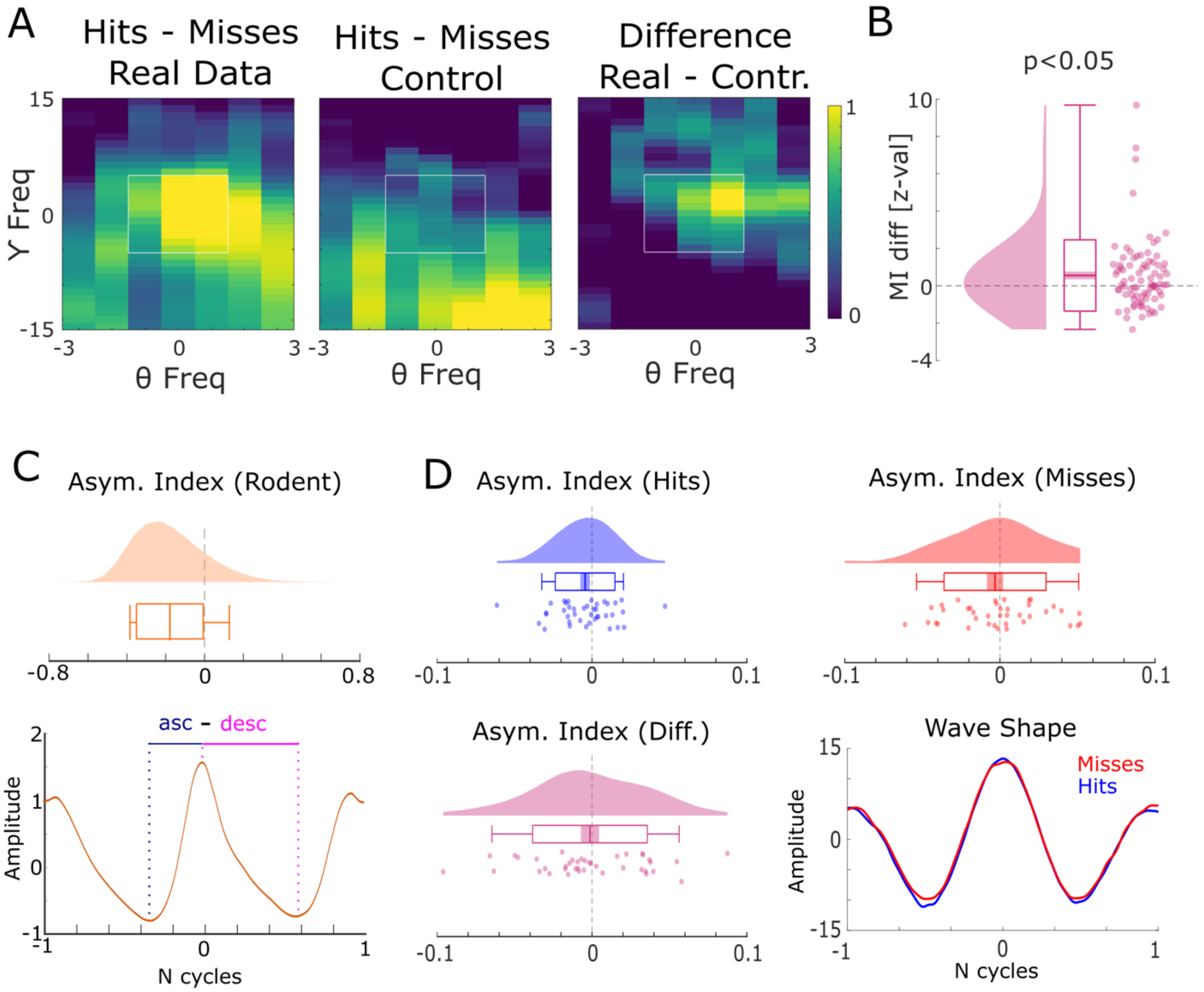


**Fig. S10. Harmonic and asymmetric waveshape control analysis**

A) The results of a harmonic control analysis are shown where the gamma power providing frequency was taken as the 8th harmonic of the theta phase providing frequency. The panel on the left shows the modulation index (MI) of the real data, the middle panel plots the MI from the harmonic control, and the panel on the right shows the difference. The white square highlights the window that was used for statistical analysis shown in (B). CFC for the real data is stronger than in the harmonic control data, ruling out an influence of asymmetric theta waveshapes. (C) Theta waveshape was quantified by the asymmetry index (see *(13)*) and is shown for a rodent LFP dataset recorded in the entorhinal cortex during an open field navigation task (courtesy of Ehren Newman). A strong asymmetry is present with the ascending flank covering less time than the descending flank. (D) The results of same analysis are shown for the theta providing channels in humans for hits (blue) and misses (red). Waveshapes appear much more symmetric compared to rodents (note the difference in scale on the x-axis between C and D), albeit both exhibit a slight asymmetry in the same direction as observed in rodents (ascending < descending flank). Importantly, no difference between hits and misses was observed in waveshape asymmetry, thus further ruling out an influence of asymmetric theta waveshape on the observed cross-frequency coupling results.

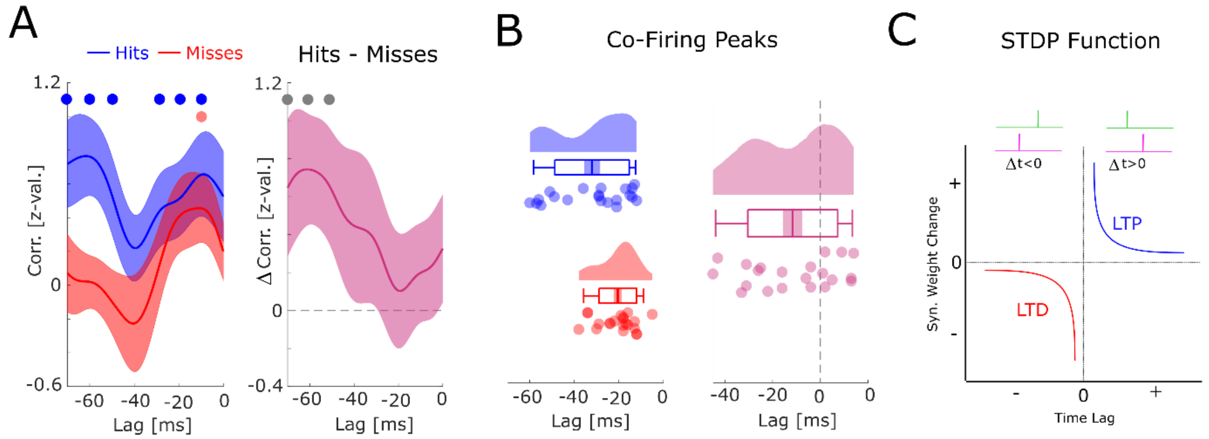


**Fig. S11. Co-firing analysis at negative lags.**

A) The results for the co-firing analysis at negative lags (putative down-stream neuron fires before putative up-stream neuron) are shown. Hits show significant above chance co-firings at lags -70 to -50, and -30 to -10 ms ($p_{corr}<0.05$), whereas misses show strongest co-firing only at lag -10 ms ($p_{uncorr}<0.05$). The co-firing difference between hits and misses is shown on the right, showing strongest differences at lags -60 to -40 ms ($p_{uncorr}<0.05$). B) Results of the co-firing peak detection analysis are plotted. Compared to hits, misses exhibit peak co-firings at shorter negative latencies (i.e. closer to 0; $t_{20}$=-2.82; p<0.05). C) A schematic of the STDP function is shown. Notably, the results shown in A and B, suggest that hits showed a tendency for putative down-stream neurons (pink) to fire *long* before putative up-stream neurons (green), whereas misses showed a tendency towards co-firing at a much *shorter* lag (-10 ms). This latter pattern would lead to a strong punishment of the synaptic connections (LTD), and hence weaker memories. Therefore, these results, albeit statistically weaker than the results for positive lags reported in the main manuscript (Figure 5), are fully consistent with STDP.

**Supplementary Table S1: Regions where distally and locally coupled neurons were recorded**

| Patient IDs | Distally coupled (up-stream) | Locally coupled (down-stream) |
|---|---|---|
| P04 | left mid. Hipp. MW4 N1 SU | left EC MW3 N1 MU |
| P04 | left EC MW8 N1 SU | left mid. Hipp. MW4 N1 SU |
| P04 | left mid. Hipp. MW7 N1 MU | left EC MW3 N1 MU |
| P05 | left mid. Hipp. MW1 N1 MU | right mid. Hipp. MW3 N1 MU |
| P05 | left mid. Hipp. MW5 N2 MU | right mid. Hipp. MW3 N1 MU |
| P07 | left mid. Hipp. MW8 N1 SU | right ant. Hipp. MW6 N1 MU |
| P07 | right mid. Hipp. MW4 N1 SU | right ant. Hipp. MW6 N1 MU |
| P07 | right ant. Hipp. MW4 N1 MU | right mid. Hipp. MW2 N2 MU |
| P07 | right ant. Hipp. MW7 N1 MU | right mid. Hipp. MW2 N2 MU |
| P07 | right mid. Hipp. MW3 N2 SU | right ant. Hipp. MW7 N2 SU |
| P07 | right mid. Hipp. MW4 N1 MU | right ant. Hipp. MW7 N2 SU |
| P08 | left ant. Hipp. MW1 N1 MU | right mid. Hipp. MW8 N1 SU |
| P08 | left ant. Hipp. MW3 N1 MU | right mid. Hipp. MW8 N1 SU |
| P08 | left ant. Hipp. MW5 N1 SU | right mid. Hipp. MW8 N1 SU |
| P08 | left ant. Hipp. MW8 N1 MU | right mid. Hipp. MW8 N1 SU |
| P08 | left ant. Hipp. MW3 N1 MU | right mid. Hipp. MW2 N1 SU |
| P08 | left ant. Hipp. MW3 N1 MU | right mid. Hipp. MW1 N1 MU |
| P08 | right mid. Hipp. MW1 N1 MU | left ant. Hipp. MW3 N1 MU |
| P08 | right mid. Hipp. MW1 N1 MU | left ant. Hipp. MW8 N1 MU |
| P09 | left ant. Hipp. MW7 N1 MU | right mid. Hipp. MW6 N1 SU |
| P09 | left PHC MW6 N1 MU | right mid. Hipp. MW6 N1 SU |
| P09 | right post. Hipp. MW2 N1 MU | right mid. Hipp. MW6 N1 SU |
| P09 | right post. Hipp. MW5 N1 MU | right mid. Hipp. MW6 N1 SU |
| P09 | left PHC MW5 N1 MU | right mid. Hipp. MW4 N1 MU |

EC = entorhinal Cortex; ant. = anterior; mid = middle; post. = posterior; Hipp. = Hippocampus; PHC = parahippocampal cortex; MW = Microwire on which neuron was recorded; N = cluster ID as obtained by WaveClus ; SU = Single Unit; MU=Multi Unit
